# Steroid sulfatase deficiency increases the frequency and persistence of ventricular arrhythmias in mice

**DOI:** 10.64898/2026.07.22.739997

**Authors:** Freya Garland-Thomas, Ailsa Saverse, Ewan D. Fowler, William Davies

**Affiliations:** Cardiff University School of Biosciences, Cardiff, UK; University of Alabama at Birmingham, Heersink School of Medicine, Alabama, USA; Division of Psychological Medicine and Clinical Neurosciences and Centre for Neuropsychiatric Genetics and Genomics, Cardiff University School of Medicine, Cardiff, UK; Cardiff University School of Psychology, Cardiff, UK; Neuroscience and Mental Health Innovation Institute, Cardiff University, Cardiff, UK

**Author notes:** to whom correspondence should be addressed: Dr William Davies, 2^nd^Floor Hadyn Ellis Building, Maindy Road, Cardiff, CF24 4HQ UK. These authors contributed equally to this work (Ewan D. Fowler, William Davies).

**Keywords:** Arrhythmias, atrial fibrillation, steroid hormones, ventricular ectopic beats, X-linked ichthyosis, Xp22.31 deletion

## Abstract

**Background:** Emerging evidence in humans has linked Xp22.31 genetic deletions to an increased risk of stress-induced arrhythmias, and association analysis across Xp22.31 has shown enrichment for atrial fibrillation genetic risk variants around *STS* (steroid sulfatase).

**Methods:** We compared heart rhythm in homozygous ‘STS-knockout’ (STS-KO) adult mice (n=19) to wildtype (WT) mice (n=11) under baseline (Tyrode’s solution), and β-adrenergic stimulation (stress), conditions using *ex vivo* perfused-heart electrocardiography (ECG). Ventricular ectopic beats (VEBs) were manually-identified, and rhythm abnormalities were quantified using a semi-automated approach.

**Results:** At baseline, the groups displayed comparable sinus cycle length and ECG intervals; all WT hearts showed stable sinus rhythm, whereas ∼30% of STS-KO hearts developed spontaneous VEBs. WT hearts typically maintained steady rhythm under β-adrenergic stimulation; in contrast, ∼60% of STS-KO hearts displayed VEBs. Under baseline and stimulated conditions the QRS interval was greater during VEBs than normal beats in STS-KO hearts, consistent with a ventricular origin. We identified a higher frequency of any abnormal beats in STS-KO hearts than in WT hearts under baseline (1.9±0.7% vs. 0.3±0.1%, p=0.038) and stimulated (5.3±2.2% vs. 0.5±0.3%, p=0.047) conditions. Under stimulated conditions, abnormal beats only occurred singly in WT hearts, whereas in STS-KO hearts, ∼50% of the time they occurred in runs of two or more.

**Conclusion:** STS deficiency in mice predisposes to arrhythmias, and the STS-KO mouse represents a tractable model for mechanistic investigation. These data support the contention that STS activity influences arrhythmic vulnerability in humans and that *STS* should be considered for inclusion in arrhythmia-related gene panels.

---

Cardiac arrhythmias predispose to cardiac arrest, heart failure, and thromboembolic events with potentially severe neurological consequences.^1^ In humans, Xp22.31 deletions encompassing *STS* (encoding the enzyme steroid sulfatase, STS) and adjacent genes, are associated with a substantially-increased risk of a range of arrhythmias (notably atrial fibrillation/flutter, AF) affecting 10-35% of carriers; association analysis across the Xp22.31 interval has revealed a nominally-significant association between common genetic variants around *STS* and an AF diagnosis.^2,3^ STS cleaves sulfate groups from steroid hormones allowing them to act as precursors for androgens and oestrogens.^4^ We aimed to directly test whether STS deficiency increases arrhythmia risk by comparing heart rhythms in mice lacking STS constitutively to those of their wildtype (WT) littermates.

C57BL/6J-Sts^em2H^/H mice harbouring a 22bp frameshift deletion in exon 2 of the pseudoautosomal *Sts* gene were obtained from the Mary Lyon Centre at MRC Harwell, the UK node of the European Mouse Mutant Archive (EMMA) (www.infrafrontier.eu; Repository number EM:15127).^5^ Homozygous ‘knockout’ (STS-KO) mice exhibit grossly-normal health despite >95% loss of peripheral STS activity, and from 20-50 weeks of age have greater normalised heart weights than their WT counterparts.^6^ Adult (12-20 weeks) WT (n=11, 7 male, 4 female) and STS-KO (n=19, 10 male, 9 female) mice were killed by stunning and cervical dislocation, and their hearts were quickly excised and perfused on a Langendorff apparatus via the aorta with oxygenated modified Tyrode’s solution.^7^ *Ex vivo* volume-conducted electrocardiograms (ECG) were recorded at 1kHz sampling rate using Ag/AgCl electrodes in a pseudo-lead II configuration in a heated solution chamber (35°C). ECGs were recorded during sinus rhythm before, and after, β-adrenergic stimulation with isoproterenol (ISO, 1µM); this manipulation mimicked stress, the most commonly-reported precipitant of arrhythmic episodes in Xp22.31 deletion carriers.^3^ Experiments and ECG analysis were independently-analysed by two investigators blinded to genotype.

Normally-distributed data according to the Shapiro-Wilk test were analysed using Welch’s or paired t-test, otherwise non-parametric equivalents were used. Categorical data were analysed using the Lancaster mid-p method. Data are presented as mean±standard error of the mean.

STS-KO and WT mice had equivalent heart:bodyweight ratios (WT:8.6±0.5mg/g; STS-KO:8.7±0.1mg/g; p=0.85). In Tyrode’s solution, STS-KO and WT hearts had comparable sinus cycle lengths (WT:183±11ms; STS-KO:171±4ms; p=0.30), PQ intervals (WT:44.2±1.5ms; STS-KO:43.3±1.1ms; p=0.63), QRS durations (WT:22.6±1.2ms; STS-KO:22.3±1.0ms; p=0.54) and QT intervals (WT:89.1±3.6ms; STS-KO:79.8±3.1ms; p=0.06). All WT hearts showed stable sinus rhythm in Tyrode’s solution, whereas ∼30 % of STS-KO hearts developed spontaneous ventricular ectopic beats (VEB) with post-extrasystolic pauses (**Fig. 1A**).

**Figure 1.**
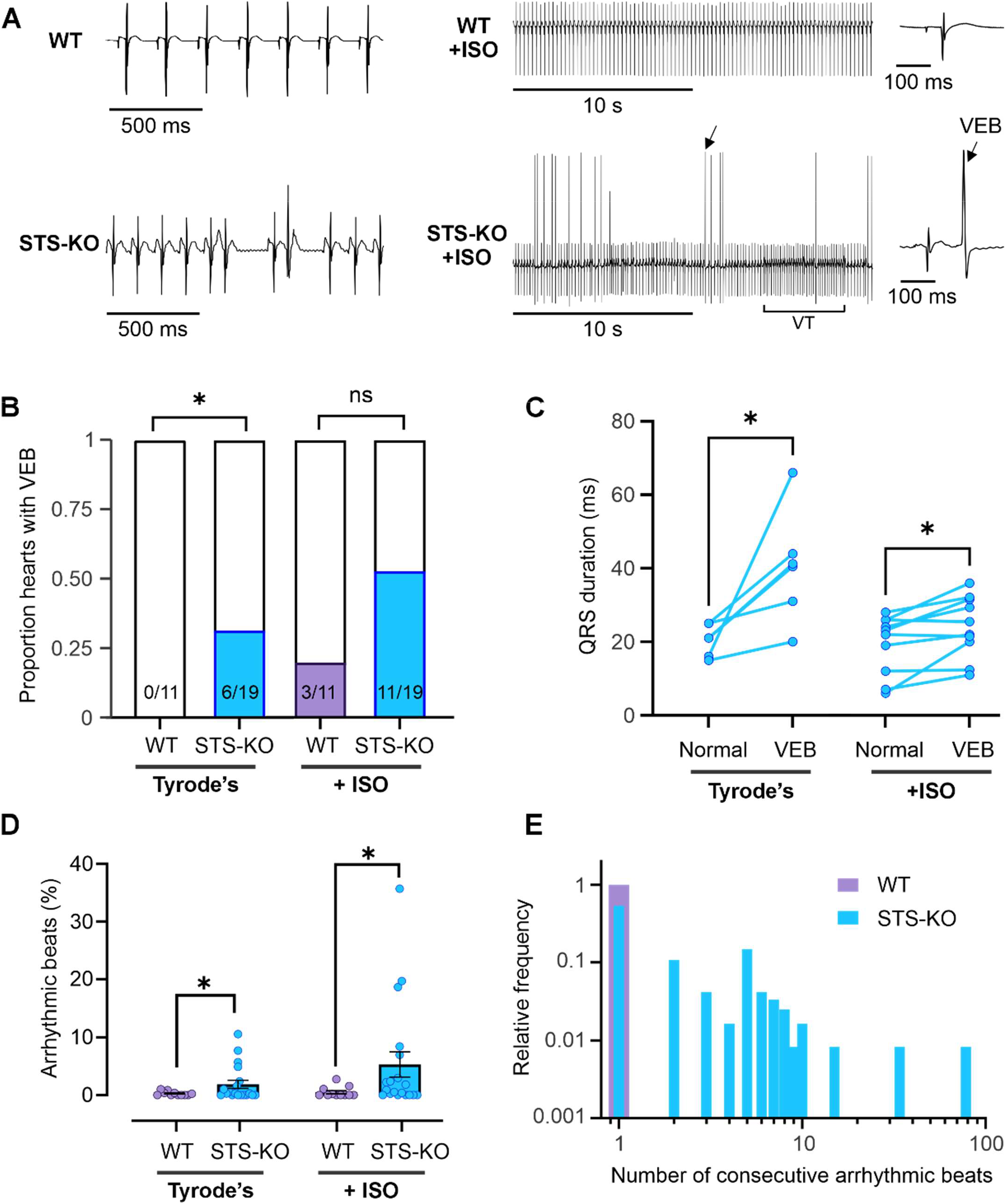
**A** Exemplar *ex vivo* ECG recorded from a WT (top) and STS-KO heart (bottom) in Tyrode’s solution (left) and during isoproterenol (ISO) stimulation (middle). The WT heart maintained stable sinus rhythm in both Tyrode’s and ISO solutions, whereas the STS-KO heart exhibited occasional ventricular ectopic beats (VEB) in Tyrode’s, which became more numerous during ISO stimulation (arrow). The inset panels (right) show the VEB indicated by the arrow on an expanded timescale. A run of spontaneous ventricular tachycardia (VT) (∼8Hz) also occurs in the STS-KO heart. **B** Proportion of hearts that exhibited VEB was greater in STS-KO than in WT in Tyrode’s solution. ISO increased the proportion of hearts with VEB in both WT and STS-KO. *p<0.05 vs WT. Number of hearts are indicated in the bars. **C** The QRS duration was greater during VEB compared to normal sinus ECG cycles from the same STS-KO hearts in both Tyrode’s (n=6 hearts) and ISO solutions (n=11 hearts). *p<0.05 vs normal QRS duration, paired t-test. **D** Cross-correlation analysis detected more arrhythmic beats in STS-KO than WT hearts in both Tyrode’s and ISO solutions. Symbols represent mean values from WT (n=11) and STS-KO (n=19) hearts. **E** Probability density function showing the number of consecutive arrhythmic beats detected during ISO stimulation. Arrhythmias in STS-KO hearts often occurred in longer runs of two or more consecutive abnormal beats.

WT hearts typically maintained steady rhythm in ISO, with only occasional, isolated, VEB observed in some hearts; in contrast, STS-KO hearts became increasingly unstable when challenged, with ∼60% of hearts displaying VEB (**Fig.1B**). In STS-KO hearts, QRS duration was significantly greater during VEB than normal beats, both in Tyrode’s solution and during ISO stimulation (paired t-test, p=0.030 and p=0.014 respectively, **Fig 1C**), consistent with a ventricular origin.

VEB can increase the likelihood of more severe tachyarrhythmias,^8^ and some STS-KO hearts developed sustained or non-sustained tachycardia, or intermittent bradycardia (**Fig.1A**). As the ECG morphology of these arrhythmic beats was more heterogenous, we used an unbiased cross-correlation algorithm-based approach to detect abnormal heart cycles, and to quantify heart rhythm abnormalities. First, all ECG cycles were identified by impulse detection of QRS complexes following previously described signal processing steps (filtering, differentiating, squaring and applying a moving integrator).^9^ An ensemble average ECG cycle was then constructed and used as a template for cross-correlation with the phase-shifted original ECG. The maximum cross-correlation value for each cycle was normalised by the autocorrelation of the template (autocorrelation factor, ACF) indicative of waveform similarity. Cycles with ACF≥15 standard deviations from the mean were considered abnormal.

Consistent with the manual identification method, our semi-automated strategy also detected infrequent, and isolated, arrhythmic beats in WT hearts in Tyrode’s and ISO solutions (**Fig.1D,E**). STS-KO hearts exhibited significantly more abnormal beats in Tyrode’s solution (p=0.038), an effect exacerbated by ISO application (p=0.047) (**Fig.1D**). Strikingly, ∼50% of all arrhythmic beats occurred in consecutive runs of two or more in STS-KO hearts during ISO stimulation (**Fig.1E**). The cycle length during runs of abnormal beats in STS-KO mice was compared to the average cycle length of normal beats from the same hearts to establish whether these were predominantly tachycardia or bradycardia; however the change in cycle length was not consistent across all hearts (p=0.92). The overall cycle length in ISO was similar between genotypes (WT:136±8ms; STS-KO:137±6ms; p=0.77).

Our results strongly indicate that STS deficiency increases arrhythmia risk in Xp22.31 deletion carriers; these individuals (comprising ∼1/1000 members of the general population) could benefit from cardiac screening. Deletion carriers are primarily ascertained through the presence of the skin condition X-linked ichthyosis (XLI), or through relation to an affected individual. XLI is associated with a high VEB burden in some individuals.^10^ Our data aligns with clinical findings that boys with XLI and abnormal heart rhythms are diagnosed with tachycardia or bradycardia in similar proportions (53 and 40%, respectively),^3^ and case studies reporting shorter QT intervals and VEB in patients with Xp22.31 deletions.^11^ The causal mechanisms linking STS deficiency and arrhythmias remain to be clarified, but could involve electrical or structural remodelling of the heart. STS deficiency in mouse skin and human keratinocyte cultures perturbs calcium signalling^12^ and gene pathways affecting cardiac septal development.^4^ STS-KO mice eventually develop cardiac hypertrophy^6^ and it is possible that a high VEB burden could contribute to this remodelling.^13^ STS-KO mice represent a highly-tractable model for mechanistic investigations and for trialling potential therapies. In humans, *STS* escapes X-inactivation, and enzyme activity is higher in female than male tissues; STS activity levels could partially explain vulnerability to arrhythmias in Turner syndrome, and sex differences in arrhythmia prevalence, course, and therapeutic response.^14^ STS inhibitors are currently being developed as therapeutics for hormone-dependent conditions;^15^ heart rhythms should be closely monitored during associated trials. Finally, Xp22.31 deletions may be responsible for ∼1/300 cases of idiopathic AF in middle-aged men.^14^ The present findings support introducing *STS* into arrhythmia-related gene panels, and assessment of its presence/sequence in idiopathic arrhythmia cases.

## Acknowledgements

Mice used in this study were obtained from the Mary Lyon Centre at MRC Harwell (MLC) and the following award is acknowledged: MC_UP_2201/2. The mouse model was generated in collaboration with Genome Editing Mice for Medicine Programme at Mary Lyon Centre MRC Harwell (https://www.har.mrc.ac.uk/projects/gemm/). FGT was supported by a Wales Heart Research Institute PhD studentship. This work was supported in part by the Physician Scientist Development Fellowship and International Medical Education at the University of Alabama at Birmingham Marnix E. Heersink School of Medicine through funding provided to AS. EDF was supported by a BHF Intermediate Basic Sciences Research Fellowship (FS/IBSRF/21/25071). Additional funding was obtained from Cardiff University School of Psychology.

## Notes

### Competing Interest Statement

The authors have declared no competing interest.

